# Cytokinin- microbiome interactions regulate developmental functions

**DOI:** 10.1101/2021.08.02.454802

**Authors:** Rupali Gupta, Dorin Elkabetz, Meirav Leibman-Markus, Elie Jami, Maya Bar

**Author notes:** **Corresponding author: Dr. Maya Bar**, Dr. Maya Bar is responsible for distribution of materials integral to the findings presented in this article in accordance with the policy described in the Instructions for Authors (www.plantcell.org).

## Abstract

The interaction of plants with the complex microbial networks that inhabit them is important for plant health. While the reliance of plants on their microbial inhabitants for defense against invading pathogens is well documented, the acquisition of data concerning the relationships between plant developmental stage or aging, and microbiome assembly, is still underway. In this work, we observed developmental-age dependent changes in the phyllopshere microbiome of tomato. The plant hormone cytokinin (CK) regulates various plant growth and developmental processes. Here, we show that age-related shifts in microbiome content vary based on content of, or sensitivity to, CK. We observed a developmental age associated decline in microbial richness and diversity, accompanied by a decline in the presence of growth promoting and resistance inducing bacilli in the phyllosphere. This decline was absent from CK-rich or CK-hypersensitive genotypes. Bacillus isolates we obtained from CK rich genotypes were found to re-program the transcriptome to support morphogenesis and alter the leaf developmental program when applied to seedlings, and enhance yield and agricultural productivity when applied to mature plants. Our results support the notion that CK-dependent effects on microbiome content support developmental functions, suggesting that these are mediated by CK in part via the bacterial community.

## INTRODUCTION

The phyllosphere microbial community plays positive roles in host plant life. Disease resistance, abiotic stress tolerance, improved vigor and alterations in life cycle phenology have been documented in the presence of specific bacterial communities (Koskella, 2020; Liu et al., 2020). The plant leaf niche occupies a large surface area, and is important for the plant microbial community structure and function. The agricultural and ecological implications of plant-beneficial interactions with microbes have motivated intense investigation into the factors that shape phyllopshere microbiota (French et al., 2021). Deciphering the factors underlying the composition and dynamics of microbiome assembly is a key step towards understanding how the microbial community affects plant health and development.

In terms of diversity and richness, the phyllosphere hosts complex microbial communities that are determined by several dynamic factors, such as plant age, plant genotype, environmental variables, geographical location and agricultural practices (Vorholt, 2012; Leveau, 2019). Previous work has uncovered factors that are central in determining the composition of microbiota. In particular, plant genotype has been identified to be an important driver that influences the structure of the phyllopshere microbiome (Bodenhausen et al., 2014; Wagner et al., 2016). In addition to host genotype, geographic growth location has also been defined as a dominant factor influencing community structure. For instance, perennial plants belonging to the same species grown in different geographic locations showed surprisingly similar leaf microbial communities than different plant species grown in close proximity (Redford et al., 2010). Recently, (Li et al., 2021) found that phyllosphere specificity varied more with respect to growth stage than to genotype of *Arabidopsis thaliana*. The growth stage and genotype of *A. thaliana* are crucial in shaping phyllosphere bacterial composition, with the former being a stronger driver. Many studies on the structure of plant-associated microbial communities have shown that plants grown in sterile conditions house microbes that resemble airborne communities, while plants grown in natural conditions often have phyllosphere communities comprised of soil microbiota (Bodenhausen et al., 2014; Maignien et al., 2014). Thus, from previous studies it becomes evident that the phyllopshere microbiome structure is complex, being influenced by various dynamic factors.

Growth stage- or age dependent bacterial community shifts in the rhizosphere have been well documented (Chaparro et al., 2014; Cordovez et al., 2021). The phyllosphere microbiome also undergoes dynamic changes as plants develop and/or age, as shown in Arabidopsis, *Lactuca sativa* and *Boechera stricta* (Williams et al., 2013; Wagner et al., 2016; Berens et al., 2019) These age-related shifts in microbial content are presumably linked with age-dependent changes in the plant, such as hormonal and/ or physiological variation. Plant aging differentially affected the abundance of multiple leaf-associated microbial taxa such as Actinobacteria, Armatimonadetes and Verrucomicrobia, at various sites in *Boechera stricta* (Wagner et al., 2016). In the phyllopshere, age-related microbiome differentiation may be associated with the differences in the leaf structure or geometry, cuticle structure, trichome placement, or composition of the volatile substances secreted by the leaf. We recently reported that leaf structural niches influence phyllosphere microbial content in different genotypes (Gupta et al., 2021).

Plant age and developmental status are important factors influencing host immune responses (Develey-Rivière and Galiana, 2007; Berens et al., 2019). (Redford et al., 2010). Plants have been shown to have differential age-dependent immune responses at the organ level (Zeier, 2005). In *A. thaliana*, young rosette leaves exhibit greater SA accumulation and SA-mediated resistance than older rosette leaves (Zeier et al., 2005). Age-dependent fluctuations in host resistance can assist plants in prioritizing the protection of valuable tissues, such as young leaves (McCall and Fordyce, 2010). However, little is known about the relationships between plant growth or developmental stage, and bacterial communities in the phyllosphere.

The plant hormone cytokinin (CK) regulates various developmental processes, including embryogenesis, cell division and differentiation, shoot and root apical meristem maintenance, shoot and root lateral organ formation, and many others (Kieber and Schaller, 2018; Gupta et al., 2021). Thus, it is not surprising that changing endogenous CK content or signaling would cause alterations to plant development, resulting in changes to organ structure and patterning. CK has been demonstrated to promote morphogenesis and delay differentiation during plant development in many different plant species and developmental contexts (DeMason, 2005; Nikolić et al., 2006; Marsch-Martínez et al., 2012; Li et al., 2013; Israeli et al., 2021), likely by delaying the differentiation of meristematic cells (Bartrina et al., 2011). Tomato plants with altered CK content have altered developmental programs, and modified organ structures. Overexpressing the CK biosynthesis gene *ISOPENTENYL TRANSFERASE7 (IPT7)*, resulting in elevated endogenous levels of CK (Shani et al., 2010), or mutating in the MYB transcription factor *CLAU*, resulting in increased CK sensitivity (Bar et al., 2016), results in highly patterned and complex leaves, Concomitantly, decreasing endogenous levels of CK by overexpressing *CK OXIDASE3 (CKX3)* (Shani et al., 2010), results in simplified leaves bearing less organs (Shwartz et al., 2016).

Recently, investigating the relationship between CK and the phyllosphere microbiome, we demonstrated that CK acts as a selective force in microbiome assembly, increasing richness, and promoting the presence of Firmicutes (Gupta et al., 2021). We found CK-mediated immunity to partially depend on the microbial community. Bacilli we isolated from CK-rich or CK-hypersensitive plant genotypes, induced plant immunity, and promoted disease resistance. Using biomimetics, we found that bacilli are preferentially supported on leaves high in CK content or signaling, due to the altered leaf structures present in theses genotypes.

Following our previous study, one of the main unanswered questions that arose was, given that CK-mediated immunity is dependent in part on the microbiome, and that CK is a driving force in microbiome assembly, could CK-mediated developmental processes also be dependent on the microbiome? In the present study, we investigated developmental-age dependent changes in the microbiome. We found that age-related shifts in microbiome content vary based on CK content/ sensitivity. Bacterial isolates from CK rich genotypes were found to re-program the transcriptome to support morphogenesis and alter the developmental program when applied to seedlings, and increase yield and productivity when applied to older plants. Our results suggest that CK-dependent effects on microbiome content and assembly support developmental functions, in line with our previous reports that CK functions are mediated via the bacterial community.

## RESULTS

### Effects of plant developmental status on the phyllosphere microbiome

To examine the effect of plant developmental stage on phyllosphere composition, microbial DNA was prepared from the phyllosphere of randomly interspersed tomato (*S. lycopersicum* M82) seedlings (10 days post germination), vegetative plants (3 weeks post germination) and reproductive flowering plants (6 weeks post germination), grown in a net house in the winter of 2018. When examining community structure between the samples using weighted UniFrac distances, we observed a significant clustering of the samples based on their developmental stage, demonstrating that the distance among biological replicates is significantly smaller within groups then between groups (**Figure 1A,B**). Interestingly, distances between the samples of the same age also decreased in parallel to the increase in developmental age, with the smallest distance observed in the oldest, reproductive group. Community richness (**Figure 1C**), Shannon index (**Figure 1D**), and proportion of Firmicutes in the bacterial community (**Figure 1E**), also all decreased as developmental age increased, while the proportion of Proteobacteria in the bacterial community increased with aging (**Figure 1F**).

**Figure 1.**
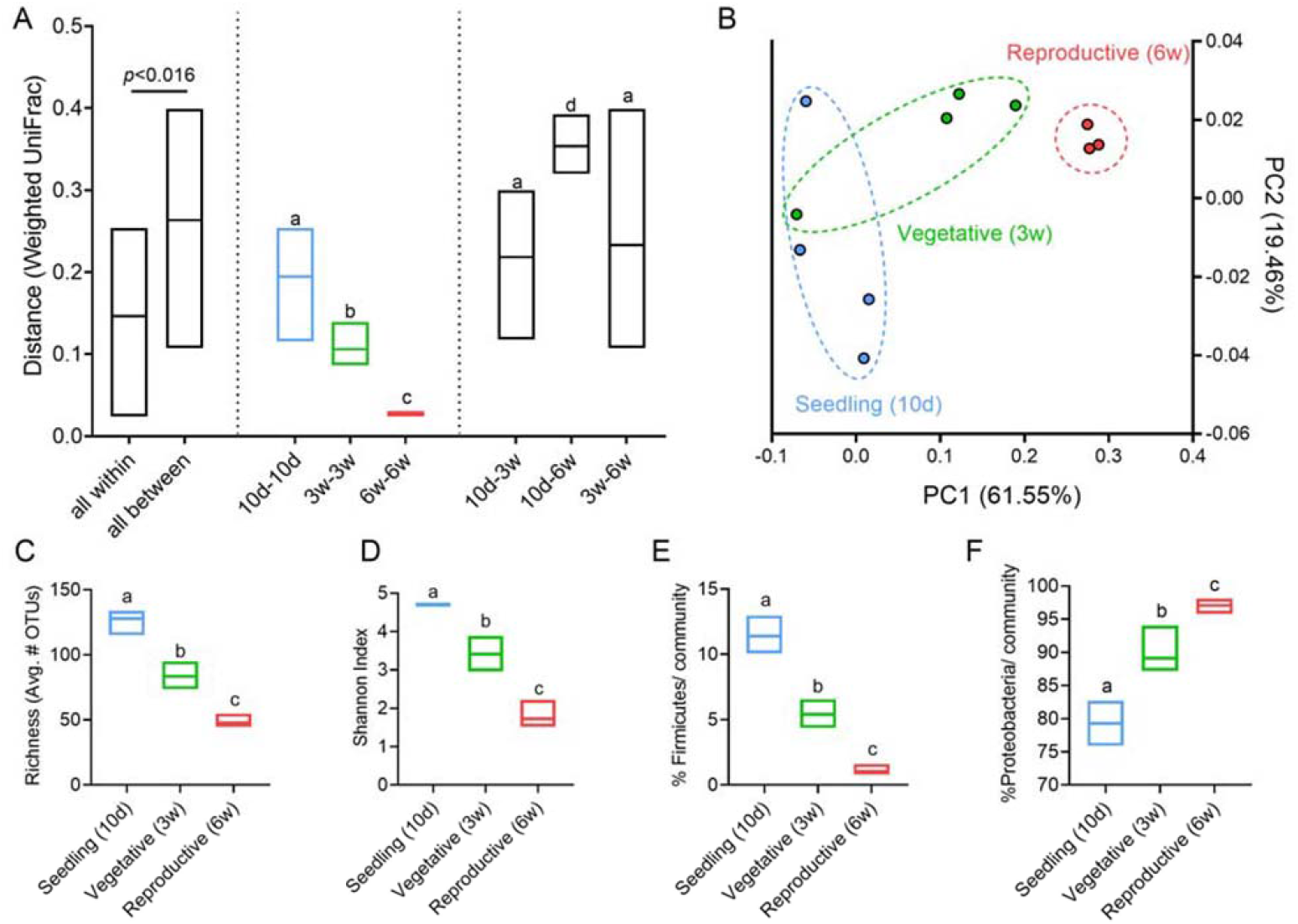
Developmental aging is accompanied by a decrease in bacterial community diversity, richness, and firmicute content. 16S rRNA sequencing of the bacterial phyllosphere of randomly interspersed S. *lycopersicum* M82 plants grown in a net house in the winter of 2018, N =4 for each genotype, of different ages: “Seedling” (10 days old), “Vegetative” (3 weeks old), and “Reproductive” (6 weeks old). **A** Weighted UniFrac beta diversity. Distance is significantly smaller within groups then between groups (p<0.016). **B** Principal coordinates analysis of distance between all individual samples in the weighted UniFrac beta diversity calculations. **C** Species richness-alpha diversity. **D** Shannon index. **E** Proportion of Firmicutes in the bacterial community of indicated genotypes. **F** Proportion of Proteobacteria in the bacterial community of indicated genotypes. Floating bars encompass minimum to maximum values, line indicates mean. Different letters indicate statistical significance between samples in a two-tailed t-test with Welch’s correction. C *p*<0.0073; D *p*<0.04; E *p*<0.0045; F*p*<0.0007.

### The amount of bacilli in the bacterial community changes throughout development in a CK dependent manner

We previously demonstrated that high CK content, or increased CK sensitivity, support an increase in phyllosphere community richness, Shannon index, and in the proportion of Firmicutes (Gupta et al., 2021). Generally, CKs are thought to be synthesized mainly in the roots and transported via the xylem to the shoots, where they exert developmental functions (Davey and van Staden, 1976; Kaminek et al., 1997). We hypothesized that the increased numbers of bacilli present in the bacterial community in seedlings (**Figure 1**) may be supported by the increased levels of CKs present in young leaves, levels which decline over time (Nordstrom et al., 2004), in accordance with the age-related decrease in Firmicutes we observed (**Figure 1E**). We examined this by assaying the amount of bacilli present in the bacterial community in seedlings and mature plants, in M82 and high and low CK content genotypes, overexpressing *pBLS≫IPT* or *pFIL≫CKX,* as well as in the high CK sensitivity mutant *clausa*. As shown in **Figure 2**, while bacilli decrease with developmental aging in M82, in the altered CK genotypes, bacilli percentage in the microbial community does not change with age. *pFIL≫IPT* and *clausa* have increased percentages of bacilli in the microbial community in both the seedling (**Figure 2A, B**) and mature (Gupta et al., 2021), **Figure 2A,C**) stages, and, unlike in the background M82 (**Figures 1, 2A**), the proportion of bacilli in the bacterial community does not decrease in mature plants when compared with seedlings. In accordance with our previous results (Gupta et al., 2021), we also observed higher amounts of microbial DNA in *pBLS≫IPT*, and lower amounts in *pFIL≫CKX*, suggesting that these genotypes also support increased or decreased amount of bacteria in general, respectively (**Figure S1**).

**Figure 2.**
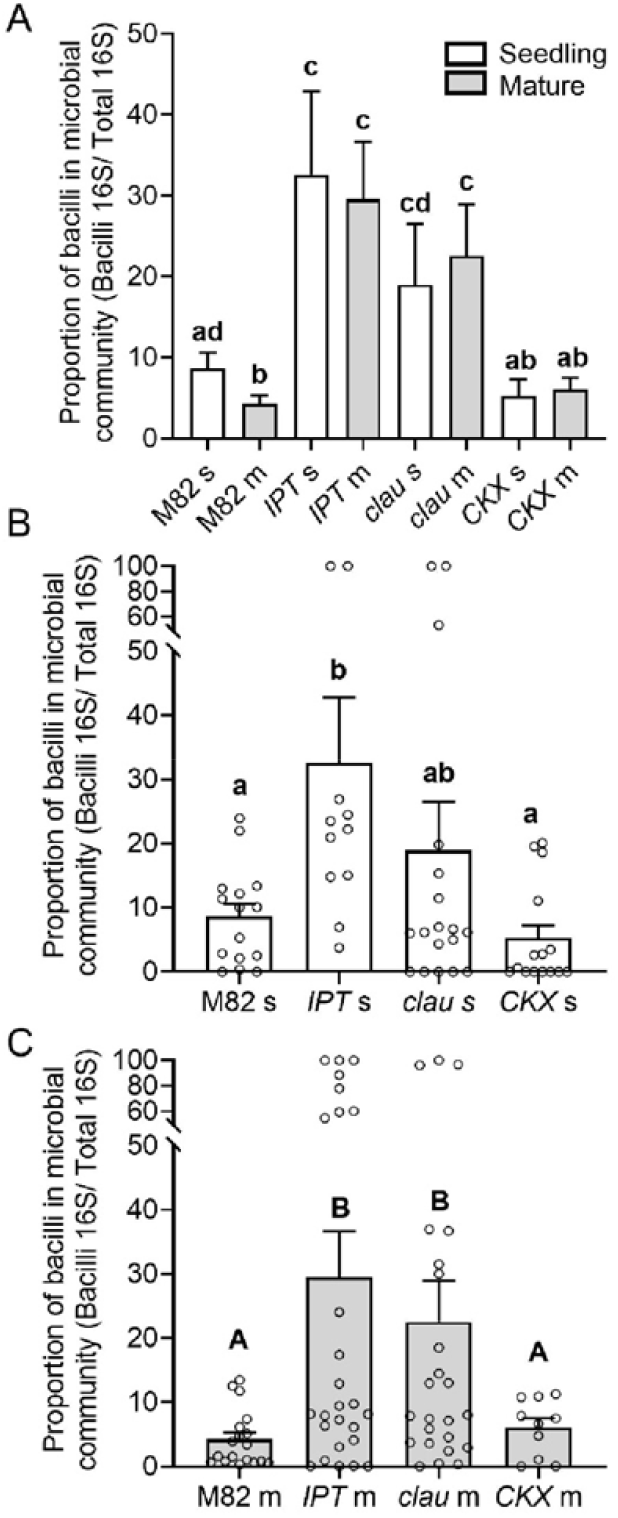
CK prevents age-associated decrease in bacilli content in the bacterial community. Bacterial DNA was extracted from indicated genotypes at the seedling (15-day old, indicated with “s” and white bars) and mature plant (45-day old, indicated with “m” and gray bars) stages. Bacilli amounts were estimated by qPCR of the bacillus 16S rRNA gene, normalized to qPCR of the total 16S rRNA gene content. (A) Comparison of bacilli content in the background M82, the high CK content I PT, the high CK sensitivity *clau*, and the low CK content CKX, at both seedling and mature stages. (B) Comparison of the different genotypes at the seedling stage, all points shown. (C) Comparison of the different genotypes at the mature stage, all points shown. Graphs depict mean ±SE. Different letters indicate statistically significant differences in an unpaired two-tailed t-test with Welch’s correction, N=10. (A) *p*<0.048. (B) *p*<0.043. (C) *p*< 0.018.

### Phylloshpere isolated bacilli from high-CK genotypes accelerate development

Bacilli are well known to have growth-promoting effects (Miljaković et al., 2020). In our previous work, we characterized several bacilli isolates from different species obtained from the phyllosphere of *pBLS≫IPT*, a high CK content genotype, finding them to promote plant immunity and disease resistance (Gupta et al., 2021). Given that seedlings, which are more morphogenetic, rapidly generating new organs, and growing at a faster pace than mature plants, have more CK and support more bacilli, we next investigated whether our phyllosphere bacilli isolates could affect seedling development. We examined the early development of tomato seedlings following treatment with different bacterial isolates. We found that two bacterial treatments, one at cotyledon emergence, and the second after one week, were sufficient to induce accelerated growth and generation of leaves in the treated seedlings, in the case of the two bacilli isolates R2E and 4C (**Figure 3A-C**). This treatment regimen also had a negative effect on growth in the case of the *Ralstonia* isolate R3C (**Figure 3A**). Differentiation of the SAM to floral meristem and sympodial meristem was also significantly increased with 4C treatment (**Figure S2**).

**Figure 3.**
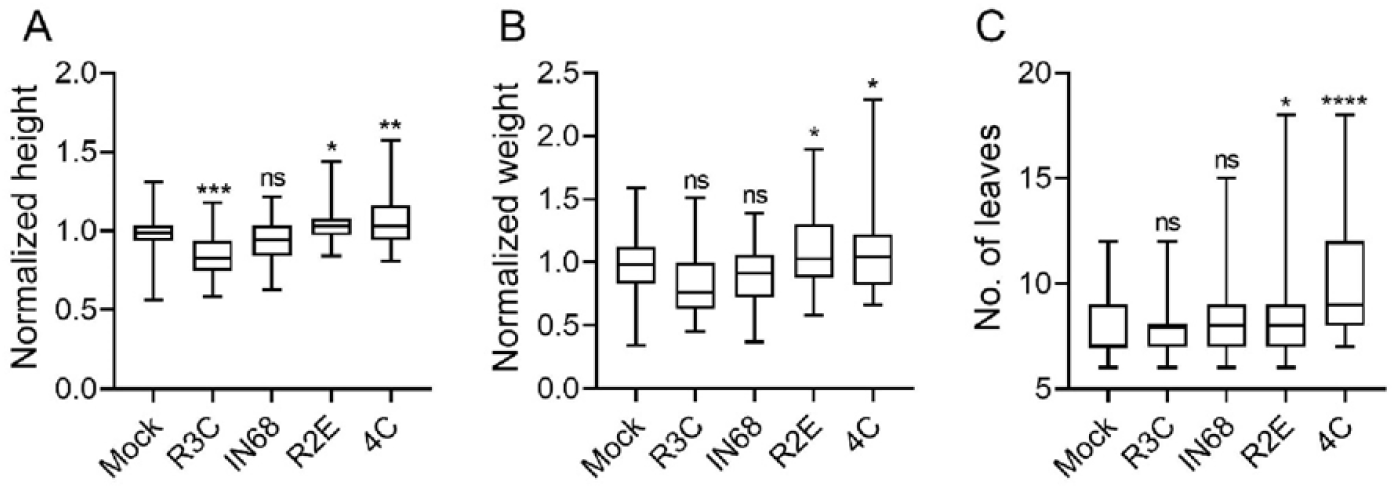
Bacilli isolated from high-CK genotypes affect development in seedlings. *S. lycopersicum* cv. M82 seedlings were treated with indicated bacteria (OD_600_=0.1), once a week for 2 weeks, starting from cotyledon emergence. Developmental parameters were measured in 10 day old M82 mock and bacterial isolate treated seedlings. (A) Seedling height (root crown to main shoot apical meristem) in centimeters. (B) Seedling weight. (C) Number of leaves produced starting from P1 (all initiated leaves were counted by dissecting the shoots). Boxplots depict minimum to maximum values, with box indicating inner quartile ranges and whiskers representing outer quartile ranges. Lines in box indicates median. Five independent experiments were conducted. Asterisks represent statistical significance from mock treatment in a one-way ANOVA with a Tukey post-hoc test (A-B), or a two-tailed t-test with Welch’s correction (C). \**p*<0.05, \*\**p*<0.01, \*\*\**p*<0.001, \*\*\*\**p*<0.0001, ns=non significant. **A** N=65, *p*<0.041. **B** N=50, *p*<0.047. C N=70, *p*<0.029.

In addition to the growth and organ initiation promoting effects, we observed (**Figure 3**) changes to the plant developmental program following bacterial treatment were an intriguing possibility. Treatment with bacterial isolates increased the number of leaves produced (**Figure 3C**). We therefore chose to examine leaf development, which follows a predictable and well characterized program *in S. lycopersicum* M82 (Shani et al., 2010; Israeli et al., 2021), in depth. For this, we selected the *B. megaterium* isolate 4C, which consistently performed best in growth promotion assays we conducted (**Figure 3**). We conducted an in depth analysis of leaf complexity, starting from the third leaf primordium (p3), in mock plants and plants treated with 4C. P3 was chosen since the first and second primordia are completely un-patterned in M82 (Bar et al., 2016). We found that starting from p3, *B. megaterium* 4C treatment results in a significant increase in leaf patterning (**Figure 4A-C**). Leaf complexity over time was also examined upon *B. megaterium* 4C or *B. pumilus* R2E treatment in seedlings (**Figure S3**). We found that both R2E and 4C increase leaf complexity (**Figure S3**), with 4C doing so earlier. Leaf complexity significantly increased during development in the mock treated plants when comparing the first and last time points among mock treatments, as expected, reflecting the “normal” developmental program.

**Figure 4.**
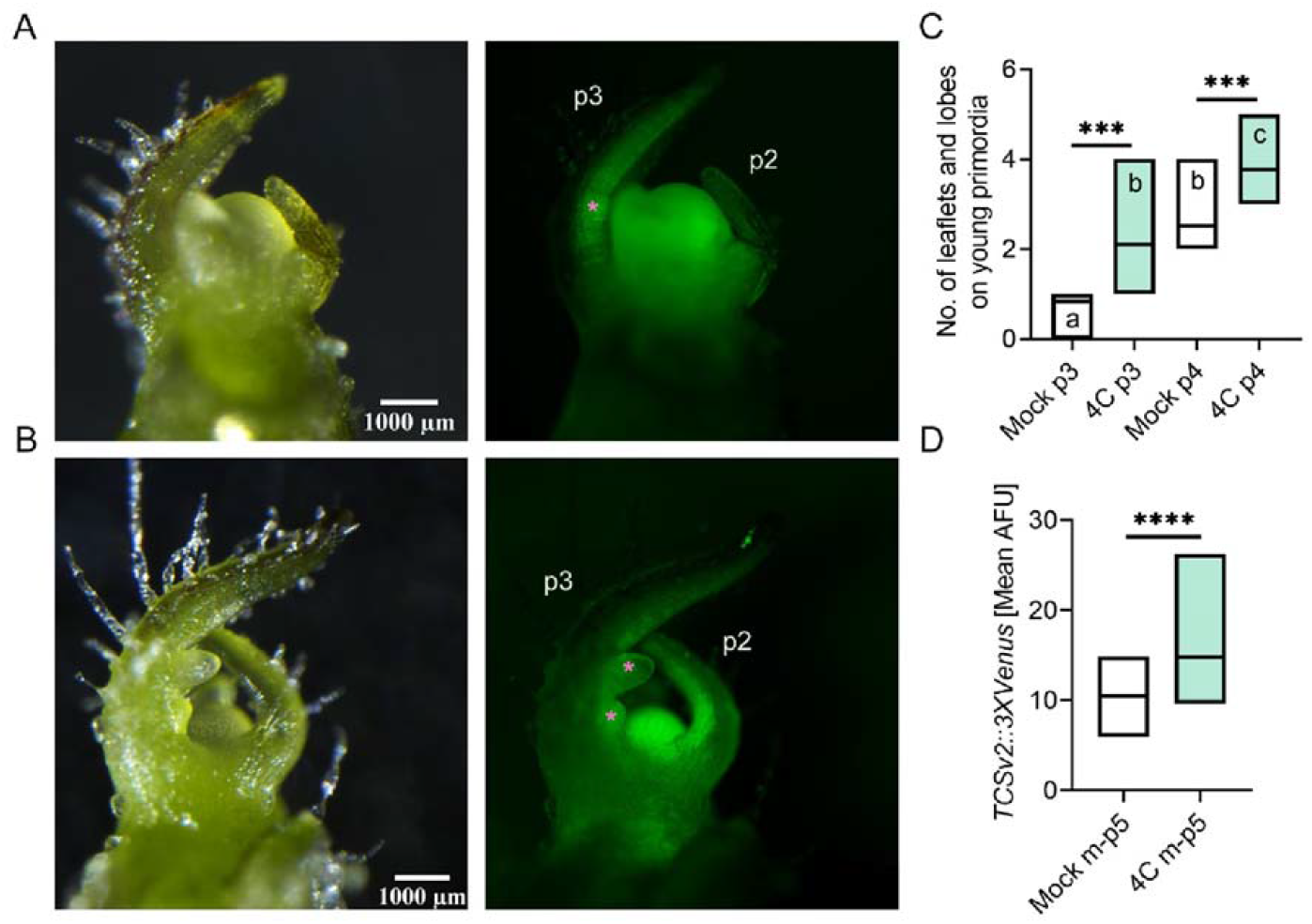
*B. megaterium* 4C accelerates leaf development and increases CK pathway activation. *S. lycopersicum* cv. M82 seedlings were treated with indicated bacteria (OD_600_=0.1), once a week for 2 weeks, starting from cotyledon emergence. Leaf complexity and TCSv2::3XVenus expression were measured in 10 day old M82 mock and 4C treated plants. (A) Typical Mock treated shoot apical meristem (SAM) and three youngest leaf primordia (p1-p3). Bar=1000 μm. (B) Typical *B. megaterium* 4C treated SAM and p1-p3. P2 and p3 are indicated in the Venus fluorescence images, with asterisks indicating the nascent leaflets on p3. (C) Number of leaflets and lobes produced on p3 and p4. (0) TCSv2 driven total Venus fluorescence was measured as mean arbitrary fluorescent units (AFU) in images captured under identical conditions in shoots comprising the 5 youngest primordia. Each primordia was quantified for leaflet number and TCS expression when Leaf No. 5 was at that developmental stage-all quantifications were done on the fifth leaf as it developed. C-D Floating bars depict minimum to maximum values, with lines indicating mean. Three independent experiments were conducted. Asterisks represent statistical significance from mock treatment, and different letters represent statistically significant differences among samples, in a one-way ANOVA with a Dunnett post-hoc test (C), or in a two-tailed t-test (D). \*\*\**p*<0.001, \*\*\*\**p*<0.0001. **C** N=12, *p*<0.0002. **D** N=21, *p*<0.0001.

### *Bacillus* treatment activates the CK response machinery and developmental genes

Using the CK activity response synthetic promoter TCS (two-component signaling sensor) fused to the VENUS fluorescent protein as a reporter (Zürcher et al., 2013), we determined that the CK pathway is activated following bacterial treatment. When examining expression of the synthetic CK-responsive promoter *TCSv2* driving Venus in transgenic M82 tomato plants stably expressing *pTCSv2::3 × VENUS*, we observed a significant increase in CK responsiveness of the leaf tissue in seedlings treated with *B. megaterium* 4C (**Figure 4A-B,D**), indicating that the accelerated development correlates with an increase in CK pathway activation.

To further characterize the effect of bacterial isolates on development, we next examined the expression of a variety of developmental genes. We chose genes related to boundary definition, which is important for organ initiation (Steiner et al., 2020; Bar et al., 2016), meristem maintenance, which is important for increased morphogenesis (Israeli et al., 2021), and, given the *TCSv2* activation results, genes of the CK pathway. We found that the bacilli isolates exclusively activated CK pathway genes (**Figure 5A-E**), with *B. megaterium* 4C not surprisingly activating more CK pathway genes than *B. pumilus* R2E. The expression of CK-responsive type-A tomato response regulators (*TRR*s) increased and the expression of *CKX* genes was also significantly altered (**Figure 5A,B,C**). All isolates activated the meristem maintenance KNOTTED1-LIKE HOMEOBOX (KNOXI/*TKN2*) (**Figure 5F**) (Avivi et al., 2000; Shani et al., 2009), while only the bacilli activated the differentiation MYB transcription factor *CLAU* (**Figure 5G**) (Bar et al., 2016), and the organ boundary determination CUC transcription factor *GOB* (**Figure 5H**) (Bar et al., 2015). To verify the response to bacterial treatment, we examined SA pathway activation using *PR1a* (**Figure 5I**) (Gupta et al., 2020), and JA pathway activation using *LoxD* (**Figure 5J**) (Dimopoulou et al., 2019). We found that all isolates activated the SA pathway (**Figure 5I**), however, interestingly, only the bacilli isolates activated the JA pathway (**Figure 5J**).

**Figure 5.**
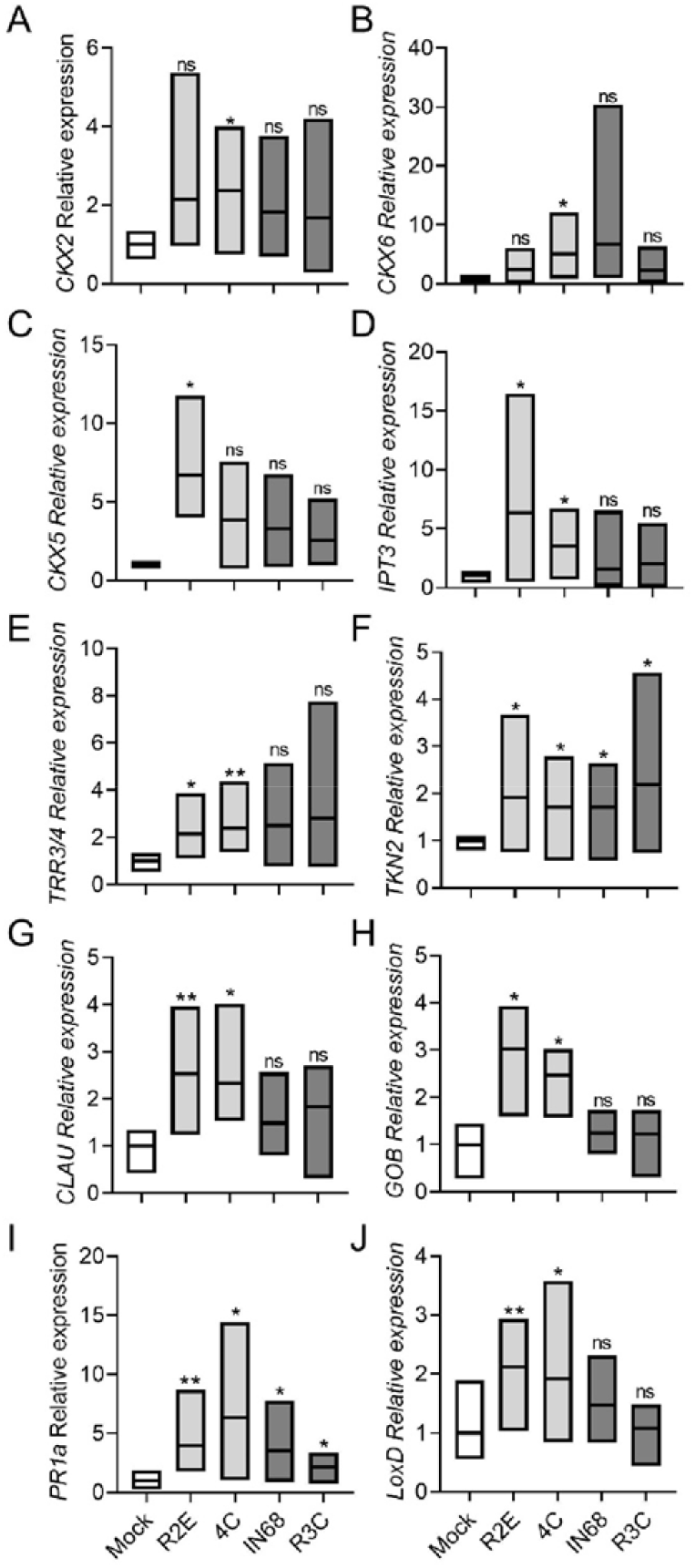
Bacilli from high-CK genotypes differentially activate morphogenetic genes and CK pathway genes. *S. lycopersicum* cv. M82 plants were pre-treated with indicated bacteria (OD_600_=0.1) once a week, two treatments in total, starting from cotyledon emergence. Gene expression was assayed by qRT-PCR, 3 days after the second treatment. Floating bars indicate minimum to maximum values (box) with mean (line in box). Bacilli are indicated in pale gray bars, gram negative bacteria in dark gray bars. Genes were normalized to a geometric mean of the expression of 3 normalizers: *SIExp, SICYP*, and *SIRPL8*. **A**- *SICKX2*; **B**- *SICKX6*; **C**- *SICKX5*; **D**- *SIIPT3*; **E**- *SITRR3/4*; **F**- *SITKN2*; **G**- *SICLAU*; **H**- *SIGOB*; **I**- *SIPRlao*, **J**- *SILoxD*. Asterisks indicate statistical significance from Mock treatment in an unpaired two-tailed t-test with Welch’s correction, N=6, *p*<0.05. (\**p* value<0.05; \*\**p* value<0.01; ns-non significant).

### Phylloshpere isolated bacilli from high-CK genotypes promote growth and increase agricultural productivity in mature plants

Since we found that bacilli isolates from high-CK genotypes accelerated development and growth in seedlings, we next examined whether they could affect growth and agricultural productivity in older plants. Several instances of agricultural use for bacilli have been reported (reviewed in (Miljaković et al., 2020). Treatment with the bacillus isolate *B. megaterium* 4C increased plant height (**Figure 6A**) and decreased apical dominance (**Figure 6B**), increased the number of leaves (**Figure 6C**), number of inflorescences (**Figure 6D**), as well as the average yield per plant (**Figure 6E**) and harvest index (**Figure 6F**). Treatment with the bacillus isolate *B. pumilus* R2E decreased apical dominance (**Figure 6B**), and increased the number of leaves (**Figure 6C**), as well as the average yield per plant (**Figure 6E**) and harvest index (**Figure 6F**), but did not affect plant height (**Figure 6A**) or the number of inflorescences (**Figure 6D**). The gram negative controls, *R. picketti* R3C and *Pseudomonas aeruginosa* IN68, had no effect on agricultural parameters, except for an increase in the number of observed inflorescences with IN68 (**Figure 6D**). A control *B. subtilis* lab strain, SD491, also increased some of the tested agricultural parameters vis., height (**Figure 6A**), number of inflorescences (**Figure 6D**), yield (**Figure 6E**), and harvest index (**Figure 6F**). None of the bacterial strains significantly affected fruit sugar content (**Figure 6G**).

**Figure 6.**
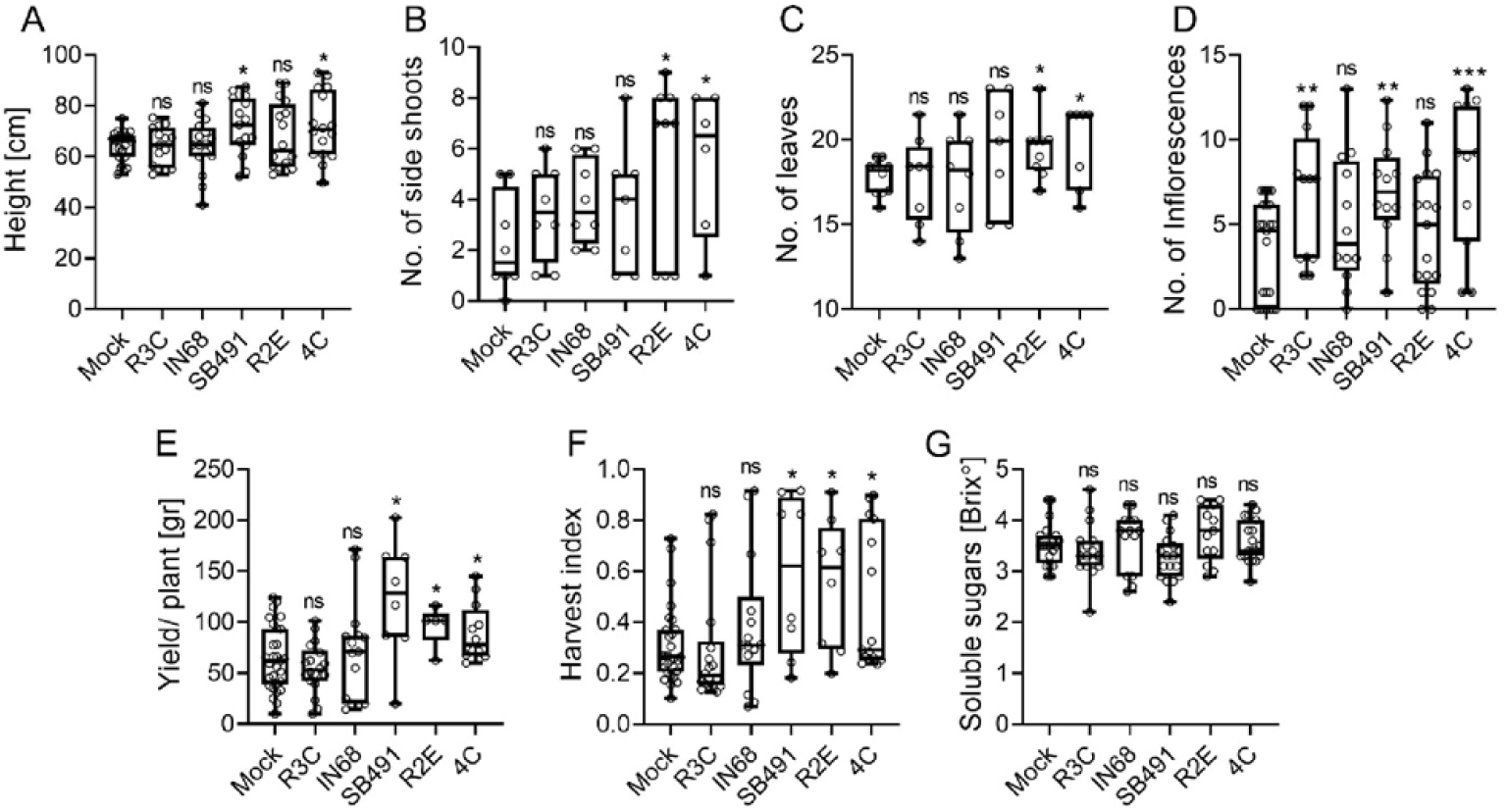
Bacilli from high-CK genotypes increase agricultural productivity. 3 week-old *S. lycopersicum* cv. M82 plants were treated with indicated bacteria (OD_600_=0.1), once a week for 4 weeks. Agricultural parameters were measured in M82 mock and bacterial isolate treated plants, just prior to harvest (65-75 days after germination). (A) Plant height (root crown to main shoot apical meristem) in centimeters. (B) Number of side shoots used as a measure of apical dominance. (C) Number of leaves produced. (D)The average number of inflorescences per plant. (E) Average yield, expressed as the total fruit weight per plant in grams. (F) Harvest index (HI), calculated as the ratio between the total mass of fruit yield and the total biomass. (G) Total soluble sugars were measured using a refractometer and are expressed as °Brix. Boxplots depict minimum to maximum values, with box indicating inner quartile ranges and whiskers representing outer quartile ranges. Lines in box indicates median. Four independent experiments were conducted. Asterisks represent statistical significance from mock treatment in a two-tailed t-test with Welch’s correction, \**p*<0.05, \*\**p*<0.01, \*\*\**p*<0.001. **A** N=16, *p*<0.035. **B** N=8, *p*<0.033. **C** N=8, *p*<0.05. **D** N=12, *p*<0.0095. **E** N=9, *p*<0.037. **F** N=9, *p*<0.043. **G** N=12, ns=non significant.

## DISCUSSION

### Developmental status/ aging influence the phyllopshere microbial community

Studies of the driving forces underlying microbial community formation have revealed that both environmental variables (reviewed in (Leveau, 2019), and host genotype and age, can be defined as the key factors driving community content and assembly (Bodenhausen et al., 2014; Li et al., 2021), depending on the context of the study. Despite recent progress in our understanding of the phyllosphere microbiome, much of the variation found in the phyllosphere remains unsolved, suggesting that the driving forces shaping microbial community structure and function have not yet been adequately defined.

Our study considered the effect of plant developmental stages on microbial community composition of the phyllosphere microbiome. Analysis of the bacterial phyllosphere community dynamics throughout plant developmental stages revealed significant changes to community richness and diversity (**Figure 1**). These results are in agreement with earlier reports concerning succession of microbial communities in the phyllosphere (Wagner et al., 2016; Manching et al., 2018; Moroenyane et al., 2021). A more detailed look at the assembled phyllopshere microbiome through developmental aging revealed that the core microbiome, composed mostly of Firmicutes and Proteobacteria, was altered in the microbial communities across development, suggesting that changes in the conditions required for survival, succession, persistence, and colonization of different microbial taxa, may occur during plant development and growth. These factors were previously reported to be important in determining phyllosphere microbial content (Maignien et al., 2014).

### Age-related changes to the microbiome are affected by CK content

CK is a central driver of development and morphogenesis. CK positively regulates cell division and proliferation in the plant shoot, driving mitosis and cytokinesis, and is involved in the organization of the shoot apical meristem (Kieber and Schaller, 2018; Yang et al., 2001). CK promotes morphogenesis by delaying the differentiation of meristematic cells (Bartrina et al., 2011). Alterations to the CK pathway in tomato result in changes in leaf phenotypes (Shwartz et al., 2016). Overexpression of the CK biosynthesis gene *AtIPT7* in tomato leaves leads to the formation of highly complex leaves, whereas overexpression of the CK oxidase/degradation gene *CKX*, results in reduced leaf complexity (Shani et al., 2010; Bar et al., 2016).The *CLAU* gene promotes an exit from morphogenesis by negatively affecting CK signaling, resulting in increased leaf complexity upon its knockout in the *clausa* mutant (Bar et al., 2016). Recently, we found that increased CK content, as in *pBLS≫IPT7*, or sensitivity, as in *clausa*, can have a strong effect on shaping the microbiome (Gupta et al., 2021). We found that high CK content or signaling increased species richness while reducing distances among samples within high-CK genotypes, resulting in dominant and consistent driving forces on bacterial community structure, and favoring Gram-positive bacteria, and bacilli in particular (Gupta et al., 2021). CK was found to shape the microbiome through both structural cues, with CK-patterned leaf structures resulting in niches that are favored by bacilli, and chemical cues, with CK promoting growth of some bacilli in vitro (Gupta et al., 2021).

Here, we analyzed the content of phyllosphere microbiota from genotypes with different CK content or sensitivity, at the seedling and mature reproductive developmental ages. We found that the abundance of total *Bacilli* was lower in WT *M82 and pFIL≫CKX* than in *pBLS≫IPT* and *clausa* at both developmental stages (**Figure 2**). The mature plant results were similar in our previous microbiome analysis of these genotypes (Gupta et al., 2021). Interestingly, examination of the microbiome shift between these two developmental stages with respect to bacilli content in high-CK and low-CK genotypes showed that, while bacilli spp. content decreases with age in the background M82, bacilli remain in high amounts in the CK-rich genotypes *pFIL≫IPT* and *clausa*, where this age-related decline in bacilli content was absent (**Figure 2A**). One possibility that arises is that the significantly lower CK content, per gram tissue, in mature plants (Davey and van Staden, 1976; Kaminek et al., 1997) underlies the age-related decline in bacilli in the microbiome in WT plants. Previous studies (Davey and van Staden, 1976) have demonstrated a close correlation between developmental changes taking place in the shoot, and the amount of CK translocated from the roots. Thus, the increased levels of CKs present in seedlings might support the increased numbers of bacilli present in the microbial community in seedlings. These increased CK levels decline over time as the plant matures (Albacete et al., 2008), which could explain the parallel decline in bacilli in the microbial community that we observed in the mature plants. This points to the possible existence of a specific microenvironment associated with CK levels, which distinctly promotes an abundance in bacilli. This microenvironment could be both structural, i.e., smaller cells and increased number of trichomes in young leaves when compared to mature (Wilkens et al., 1996; Busta et al., 2017) that create an altered physical topography available for bacterial colonization, as we previously reported in connection with high-CK genotypes (Gupta et al., 2021), or direct chemical effects stemming from the actual changes in CK levels. Of note, is that we previously found CK was able to support our phyllopshere isolates in vitro, improving their growth, biofilm formation, and swarming motility, when applied in the absence of a plant (Gupta et al., 2021). Therefore, although distinct changes in structural leaf microenvironments available for colonization between seedling leaves and mature plant leaves are highly likely, it is also possible that the abundance of CK present in younger plants acts chemically to support an increased amount of bacilli in the phyllosphere.

### *Bacilli* isolated from high-CK genotypes can alter developmental programs and increase plant productivity

The microbiome has been demonstrated to be required for achieving predictable developmental outcomes, as plants in sterile or axenic environments often grow more slowly and have altered development (Kremer, 2018; Li et al., 2020). Recently, the necessity of plant growth promoting *Bacillus* in the microbial community for disease management has been restated (Chen et al., 2020). *Bacillus* spp. are well known to have plant growth promoting activities (reviewed in Miljaković et al., 2020). We reported that bacilli we isolated from *pBLS≫IPT* enhance disease resistance by triggering plant immunity (Gupta et al., 2021). A trade-off between induced disease resistance and plant growth has been reported (Berens et al., 2019; Karasov et al., 2017). The bacilli isolates we obtained from high-CK plant genotypes protected tomato plants from disease (Gupta et al., 2021), while also supporting growth (**Figure 6**), suggesting a positive, rather than negative, correlation between growth and defense when tomato plants are treated with these bacteria. This could indicate an agricultural advantage to treatment with certain bacilli isolated at specific regimens, and will be investigated further. Our results indicate that the growth promotion exerted by bacilli strains isolated from high-CK environments are the result of alterations to developmental programs (**Figures 3-4, S2-S3**). During leaf development, the young leaf undergoes morphogenesis and reaches the mature, differentiated stage of its development simultaneously with the decline in its morphogenetic potential. Compound leaves of tomato are composed of multiple leaflets, which initiate basipetally from a meristematic region at the leaf margin known as the marginal blastozone (Shani et al., 2010; Hagemann and Gleissberg, 1996; Steiner et al., 2020). The leaf morphogenetic potential is harbored by meristematic cells, which respond to CK and therefore exhibit *TCS* activation. *TCSv2* driven expression was observed in an expanded region in *B. megaterium* 4C treated tomato seedlings in the shoot apical meristem (SAM) and three youngest leaf primordia (p1-p3), demonstrating that CK pathway activation during leaf development was increased upon *B. megaterium* treatment (**Figure 4A,B,D**). In parallel, leaves from bacillus treated plants displayed an increase in patterning, exhibiting 1-2 additional organs than typically observed on leaves of a similar developmental plastochron (**Figure 4A,B,C**), confirming that morphogenesis is indeed promoted by *B. megaterium* treatment.

CK pathway genes regulate the activity of meristems (Bartrina et al., 2011). The KNOXI gene *Tkn2*, plays important role in promoting leaf morphogenesis by delaying differentiation, preserving the meristematic identity of the leaf margin. The NAM-CUC transcription factor GOBLET determines boundaries within meristematic regions, that are necessary for organ initiation (Berger et al., 2009; Bar et al., 2016) While the MYB transcription factor *CLAU* regulates the exit from the morphogenetic phase of tomato leaf development by affecting the CK/ GA balance (Israeli et al., 2021). The changes in the expression levels of these genes upon treatment with bacilli isolated from high-CK genotypes (**Figure 5**) further supports the notion that these particular bacilli isolates boost the leaf morphogenetic potential, in part through the promotion of CK signaling. Possibly, these effects are also mediated by bacterial CK produced by these bacilli isolates for the purpose of their interaction with the host plant they colonize, though further work is needed to examine the role of bacterial CKs in this interaction and determine whether plant developmental programs can be, directly or indirectly, altered by bacterial CK.

Interestingly, age-related immunity was recently suggested to be microbiome dependent (Berens et al., 2019). This raises the attractive possibility that the effect of CK on microbial content, which depends on plant age/ developmental status, could also relate to CK-mediated immunity, i.e., CK-mediated immunity is age-dependent, or age-dependent immunity is CK-mediated, depending on the context. Further work is needed to elucidate the level of overlap between these two previously described phenomena.

Given its roles in growth and development, CK basically alters aging. In high CK content, plants become more morphogenetic, meristems are supported for longer times, senescence is delayed, and thus, “juvenility” is retained, i.e., “aging” is delayed. This delay apparently causes a lengthening of the developmental and temporal windows that support bacilli in the phyllosphere, leading to both increased growth and development (Figures 3-6, S2-S3), and improved pathogen resistance (Gupta et al., 2020, 2021).

## CONCLUSION

Analyzing developmental-age related changes in the phyllosphere microbiome, we observed a developmental age associated decline in microbial richness and diversity, accompanied by a decline in the presence of growth promoting and resistance inducing bacilli in the phyllosphere. We show that this is likely caused by the parallel decline in CK content as the plant ages. Treating WT seedlings with bacilli isolated from high-CK genotypes, resulted in significant alterations to plant development, and increased agricultural productivity. This suggests that bacterial treatments, either as single isolate or in a consortia context, could be examined in order to “re-introduce” these beneficial microbial community members that are lost during aging, or prevent their loss from occurring. Additional work is needed to examine the performance of these bacilli in agricultural settings.

## Acknowledgements

The authors wish to thank Stefan J Green and Jonathan Friedman for helpful discussions, and the Bar and Jami group members for continuous discussion and support.

## Author contributions

Conceptualization: MB. Design: MB and RG. Methodology: RG, ML-M, EJ, and MB. Experimentation: RG, DE, ML-M and MB. Analysis: RG, DE, ML-M, EJ, and MB. Manuscript: RG and MB.

## Data availability Statement

The authors declare that the data supporting the findings of this study are available within the paper and its Supplementary information files. Raw data is available through NCBI-SRA, Bioproject PRJNA729221.

## METHODS

### Plant materials and sample collection

During the winter of 2018, tomato leaf samples were collected from a roofed net house, 2 mm nylon mesh net, in ARO, Volcani Institute, Rishon Lesion, Israel., Genotypes used, all in the cv. M82 background, were as follows: M82 background line; *pBLS»IPT7*, which contains elevated endogenous levels of CK-referred to hereinafter as “*pBLS»IPT*” or “IPT”; *clausa*, which has increased CK sensitivity coupled with decreased CK content, referred to hereinafter as “*clausa*” or “*clau*”; and the CK depleted *pFIL»CKX3*, referred to hereinafter as “*pFIL»CKX*” or “CKX” (Gupta et al., 2021).

### 16S rRNA amplification, amplicon sequencing and bioinformatic analysis

To examine whether plant developmental age affects tomato phyllosphere composition, phyllosphere microbial DNA was extracted from *S. lycopersicum* cv. M82 at various developmental ages. Plants were transplanted into the nethouse at randomly interspersed locations, and all samples were collected when the latest developmental stage was reached (for the oldest plants) on the same sampling date. For phyllosphere DNA isolation, five leaflet samples per sampling age were collected from the middle lateral leaflets of leaves 5-6 of 10 different plants per sample, using ethanol-sterilized forceps. Twenty mL of 0.1 M potassium phosphate buffer at pH 8 were added to the tubes. The samples were sonicated in a water bath for 2 min and vortexed for 30 s twice. The pellet of microbes was obtained after centrifugation at 12,000 g for 20 min at 4°C. The pellet re-suspended in potassium phosphate buffer (Gupta et al., 2021). Total DNA from tomato phyllosphere microorganisms was isolated using modified protocols described by (Yang et al., 2001) and (Tian et al., 2017), and used as a template for 16S rRNA PCR amplification. 16S rRNA amplicons were generated with the following primers: CS1_515F:5’- ACACTGACGACATGGTTCTACAGTGCCAGCMGCCGCGGT-‘3;CS2_806R: 5’- TACGGTAGCAGAGACTTGGTCTGGACTACHVGGGTWTCT-’3 (Green et al., 2015). Amplicon sequencing was conducted at the UIC core facility, using Illumina MiSeq sequencing. QIIME 1.9 (Caporaso et al., 2010) was used for basic bioinformatics analysis: read merging, primer trimming, quality trimming, length trimming, chimera removal, clustering of sequences, annotation of clusters, and generation of a biological observation matrix (BIOM; sample-by-taxon abundance table). Taxonomy for the operational taxonomic units (OTUs) was assigned using BLAST against the Silva database (Glöckner et al., 2017) (silva_132_16S.97). Alpha and beta-diversity, and Shannon index, were performed with QIIME 1.9 as well the workflow script core_diversity_analysis.py. The sequence data generated in this study was deposited to the Sequence Read Archive (SRA) at NCBI under PRJNA729221.

### Quantification of leaf bacteria through DNA qPCR

Total DNA extracts were used for quantification of specific genes using qPCR. Total 16S rRNA and bacillus genes copy numbers were obtained using the 16S rRNA 515F/806R primer pair (515f: 5’- GTGCCAGCMGCCGCGGT-’3 and 806R: 5’-GGACTACHVGGGTWTCT-’3) (Green et al., 2015) and bacillus specific BacF/BacF (BacF: 5’- AGGGTCATTGGAAACTGGG-’3 and 806R: 5’-CGTGTTGTAGCCCAGGTCATA-’3) (Kuske et al., 1998), respectively. The quantification was performed with a Rotor-Gene Q machine (Qiagen) detection system and Power SYBR Green Master Mix protocol (Life Technologies, Thermo Fisher, United States). The standard regression curve was obtained using a *B. megaterium* 16S rRNA gene fragment and serial 1:10 dilutions. Four replicates of each standard dilution were prepared to generate a mean value. The standard regression curve was prepared to determine the gene copy numbers in the unknown samples, and numbers were normalized to the standard sample. All PCR reactions were performed in triplicates.

### Bacterial isolate treatments

Epiphytic bacteria were isolated and identified as described (Gupta et al., 2021). Accession numbers and details of bacterial isolates used in this study are provided in Table 1.

**Table 1-.** bacterial isolates used in this work.

| <u>ID</u> | <u>Accession</u> | <u>Species</u> | <u>Source</u> |
| --- | --- | --- | --- |
| 4C | MZ148746 | *Bacillus megaterium* | *pBLS>>IPT* isolate |
| R3C | MZ148747 | *Ralstonia pickettii* | *pBLS>>IPT* isolate |
| R2E | MZ148745 | *Bacillus pumulis* | *pBLS>>IPT* isolate |
| IN68 | -- | *Pseudomonas aeruginosa* | Wheat phyllosphere isolate obtained from____Jonathan Friedman, Hebrew University of Jerusalem |
| SB491 | SB491<br>"legacy"<br>strain (Zeigler et al., 2008; Friedman et al., 2017) | *Bacillus subtilis* | Jonathan Friedman, Hebrew University of Jerusalem |

*S. lycopersicum* cv. M82 seeds were sown after surface sterilization (with 1.5% NaOCl for five minutes, followed by three rinses with sterile water) in a tray containing potting mixture. After germination, a single tomato seedling was transplanted to each pot (0.5 L, diameter=10 cm) containing green quality soil mix, Tuff soil, Israel. Pots were kept in the nethouse at ambient temperature (Day-20°C-26°C; Night-12°C-20°C), 12-h photoperiod. Bacterial colonies of *B. pumulis* R2E, *B. megaterium* 4C, *R. pickettii* R3C, *P. aeruginosa* IN68 and *B. subtilis* SB491 from a 24 h plate culture were washed twice in sterile distilled water, and then re-suspended in a 10 mM MgCl2 solution. The cell suspension was adjusted to an optical density of OD_600_= 0.1 (approximately equal to 10^8^ CFU mL^-1^) using a spectrophotometer (Tecan). For mature plants (4-5 weeks old at the start of the experiment), soil drenching of the plants was carried out by pouring 10 mL of bacterial suspension into each pot once a week, for four weeks. For seedlings, plants were spray-drenched using a hand-held spray bottle once a week, for two weeks, starting from cotyledon emergence. Plants treated with sterile distilled water served as controls.

### Plant RNA preparation and qRT-PCR

RNA was isolated from liquid N2 ground shoot apices of 10 day old seedlings, including the shoot apical meristem (SAM) and P1-P5 leaf primordia, of 12 to 15 seedlings individually treated with bacteria viz., *B. pumulis* R2E, *B. megaterium* 4C, *R. pickettii* R3C, and *P. aeruginosa* IN68, using Tri reagent (Sigma-Aldrich) as per the manufacturer’s recommendations. RNA concentrations were quantified, and cDNA was then synthesized from 2 μg RNA in a 20 μL reaction, using both reverse transcriptase and oligo(dT) primers provided with the cDNA Synthesis kit (Promega, United States). RT-qPCR was performed according to the Power SYBR Green Master Mix protocol (Life Technologies, Thermo Fisher, United States), using a Rotor-Gene Q machine (Qiagen) detection system. Primer sequences used for the qRT-PCR analyses are detailed in Supplemental Table 1 (Gupta et al., 2020; Bar et al., 2016). Expression of all assayed genes was normalized relative to tomato a geometric mean of the copy number of the three housekeeping genes, ribosomal protein *SIRPL8* (Solyc10g006580), *Slcyclophilin* (Solyc01g111170) and *SIEXP* (Solyc07g025390) was used for normalization. All primer efficiencies were in the rage 0.98-1.03 (see supplementary Table 1). Relative expression was calculated using the copy number method for gene expression (D’haene et al., 2010).

### Seedling developmental analysis, dissection, and imaging

Seedlings were harvested from soil by cutting them at the stem base. Height from the stem base to the SAM, and weight were measured using a ruler and an analytical scale, respectively. The number of leaves was counted by dissecting the shoot under a stereomicroscope and counting all the initiated leaves, starting from P1. Differentiation of the meristem to floral and sympodial follows a predictable pattern in *S. lycopersicum* M82 (Park et al., 2012; Steiner et al., 2020), and was analyzed microscopically in dissected shoots.

Leaves are produced successively on the plant, and at a given time point each leaf is at a different developmental stage. Each leaf is thus characterized by both its position on the plant (for example, L1 is the first leaf produced and L5 is the fifth), and by its developmental stage. Thus, L5 P1 is the fifth leaf when it is at the P1 stage and has just initiated from the SAM, and it becomes L5 P2 after the next primordium initiates, and so on. For each developmental stage analyzed, the fifth leaf from at least ten different plants was analyzed for leaf complexity (the amount of leaflets). For analysis of TCSv2:3XVENUS expression, dissected whole-leaf primordia were placed into drops of water on glass microscope slides and covered with cover slips. The pattern of VENUS expression was observed with a Nikon SMZ-25 stereomicroscope equipped with a Nikon-D2 camera and NIS Elements v. 5.11 software (Steiner et al., 2020).

### Data analysis

All experimental data is presented as minimum to maximum values with median or mean, in boxplots or floating bars, or as average ±SEM, with all points displayed. For microbiome analyses, differences between two groups were analyzed for statistical significance using a Mann-Whitney test, or a two-tailed t-test, with Welch’s correction where applicable (unequal variances). Differences among three groups or more were analyzed for statistical significance with a Kruskal-Wallis ANOVA, with Dunn’s multiple comparisons post-hoc test. For all other analyses, differences between two groups were analyzed for statistical significance using a two-tailed t-test, with Welch’s correction where applicable (unequal variances), and differences among three groups or more were analyzed for statistical significance with a one-way ANOVA. Regular ANOVA was used for groups with equal variances, and Welch’s ANOVA for groups with unequal variances. When a significant result for a group in an ANOVA was returned, significance in differences between the means of different samples in the group were assessed using a post-hoc test. The Tukey test was employed for samples with equal variances when the mean of each sample was compared to the mean of every other sample. The Bonferroni test was employed for samples with equal variances when the mean of each sample was compared to the mean of a control sample. The Dunnett test was employed for samples with unequal variances. All statistical analyses were conducted using Prism^8^.

## Supplemental materials

**Figure S1.**
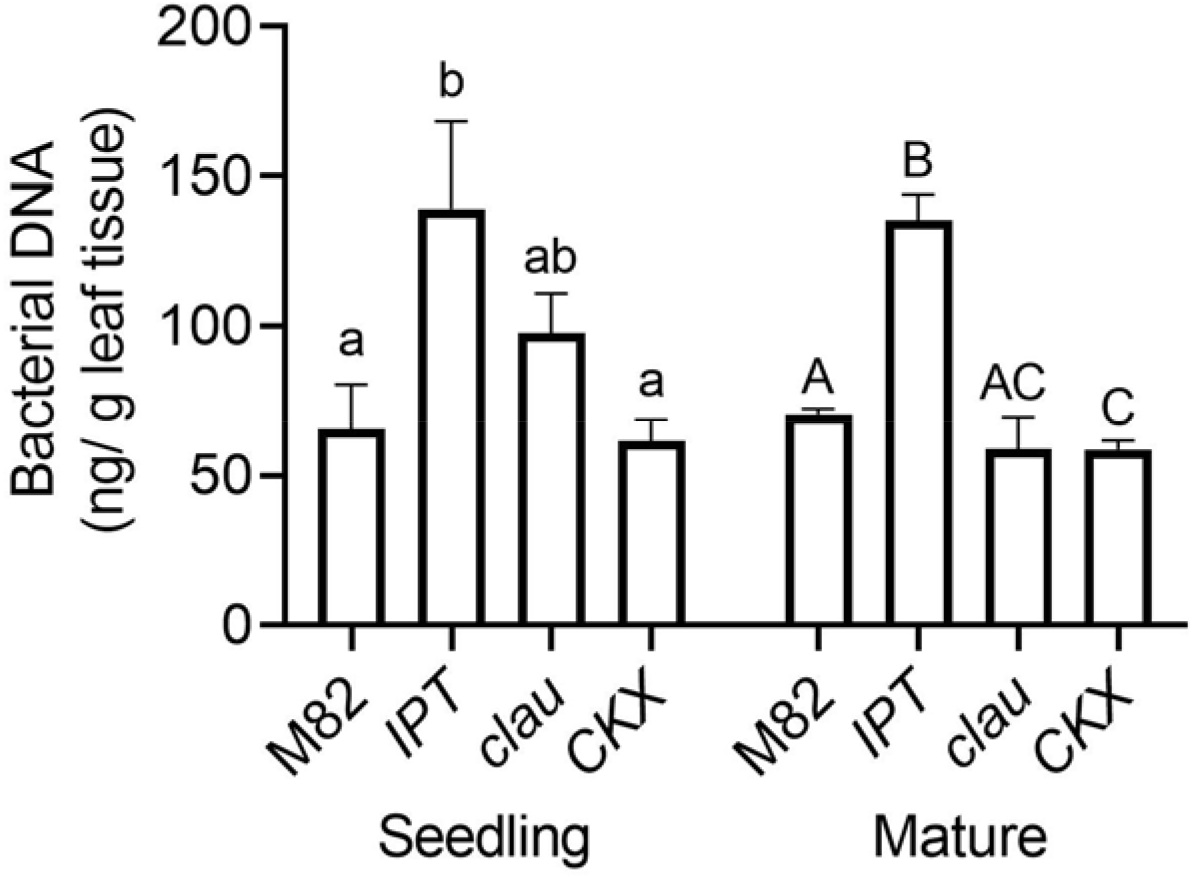
The amount of bacteria in the phyllosphere is CK dependent. Bacterial DNA was extracted from indicated genotypes at the seedling and mature plant stages. Amount of bacterial DNA obtained per gram leaf tissues is plotted. Graphs depict mean ±SE. Different letters indicate statistically significant differences in an unpaired two-tailed t-test with Welch’s correction, N=10, *p*<0.05.

**Figure S2:**
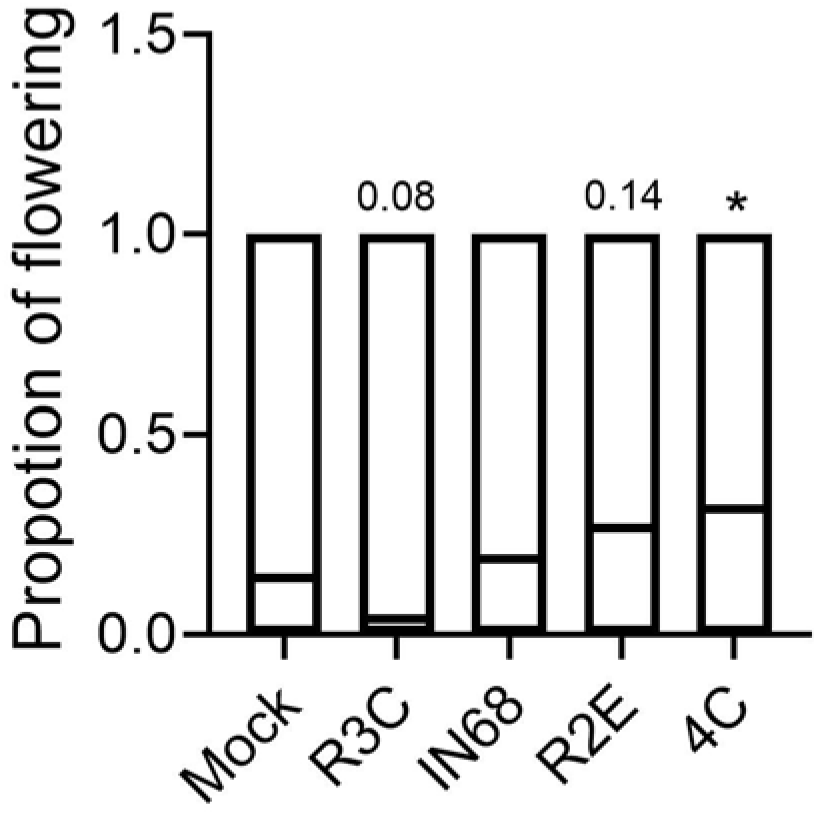
*B. megaterium* 4C induces differentiation of the flowering meristem. Presence of the floral meristem was examined in 10 day old M82 mock and bacterial isolate treated seedlings. Floating bars depict minimum to maximum values, with lines indicating mean. Five independent experiments were conducted, N=30. Asterisks represent statistical significance from mock treatment in a two-tailed t-test. \**p*<0.05.

**Figure S3:**
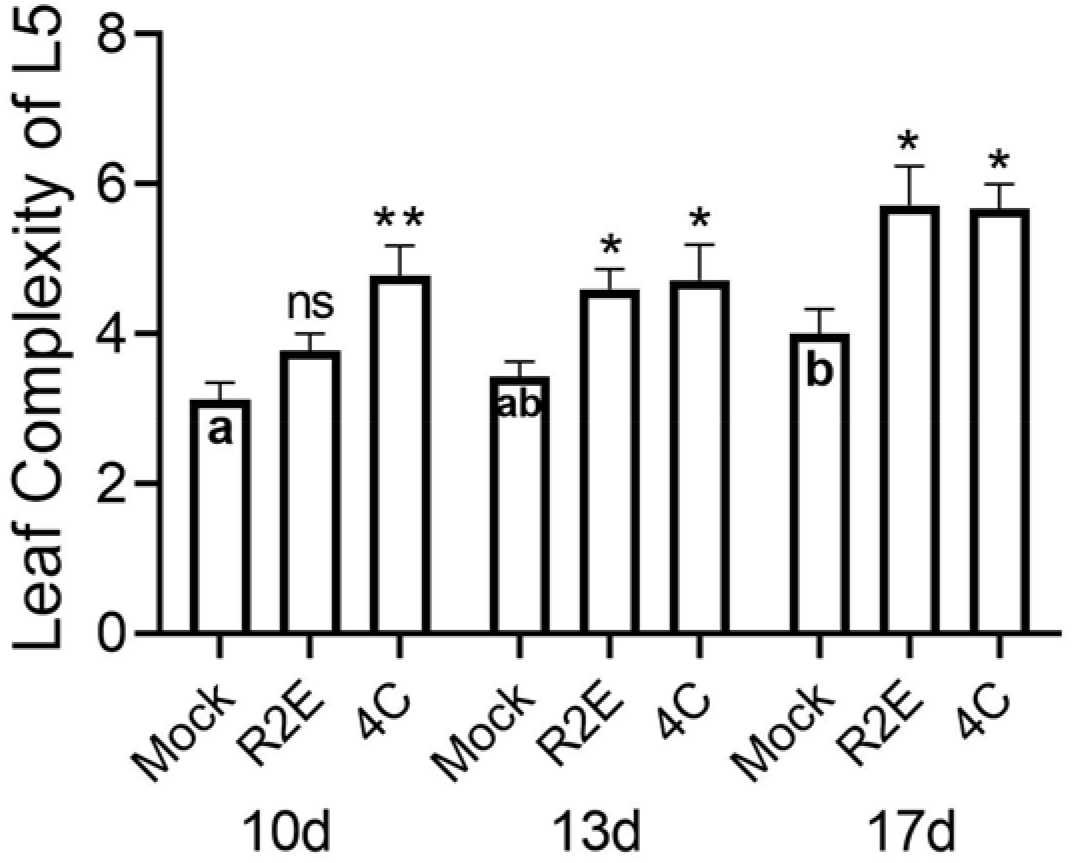
*B. megaterium* 4C and *B. pumilus* R2E accelerate leaf development-Leaf complexity over time. Leaf complexity of the fifth leaf (L5) was measured in M82 mock, R2E, and 4C treated plants over time. Bars depict mean ±SEM. Three independent experiments were conducted, N=10 for each time point. Asterisks represent statistical significance from mock treatment, and different letters represent statistically significant differences among samples, in a two-tailed t-test. \**p*<0.05, \*\**p*<0.01.

**Supplemental Table1.**
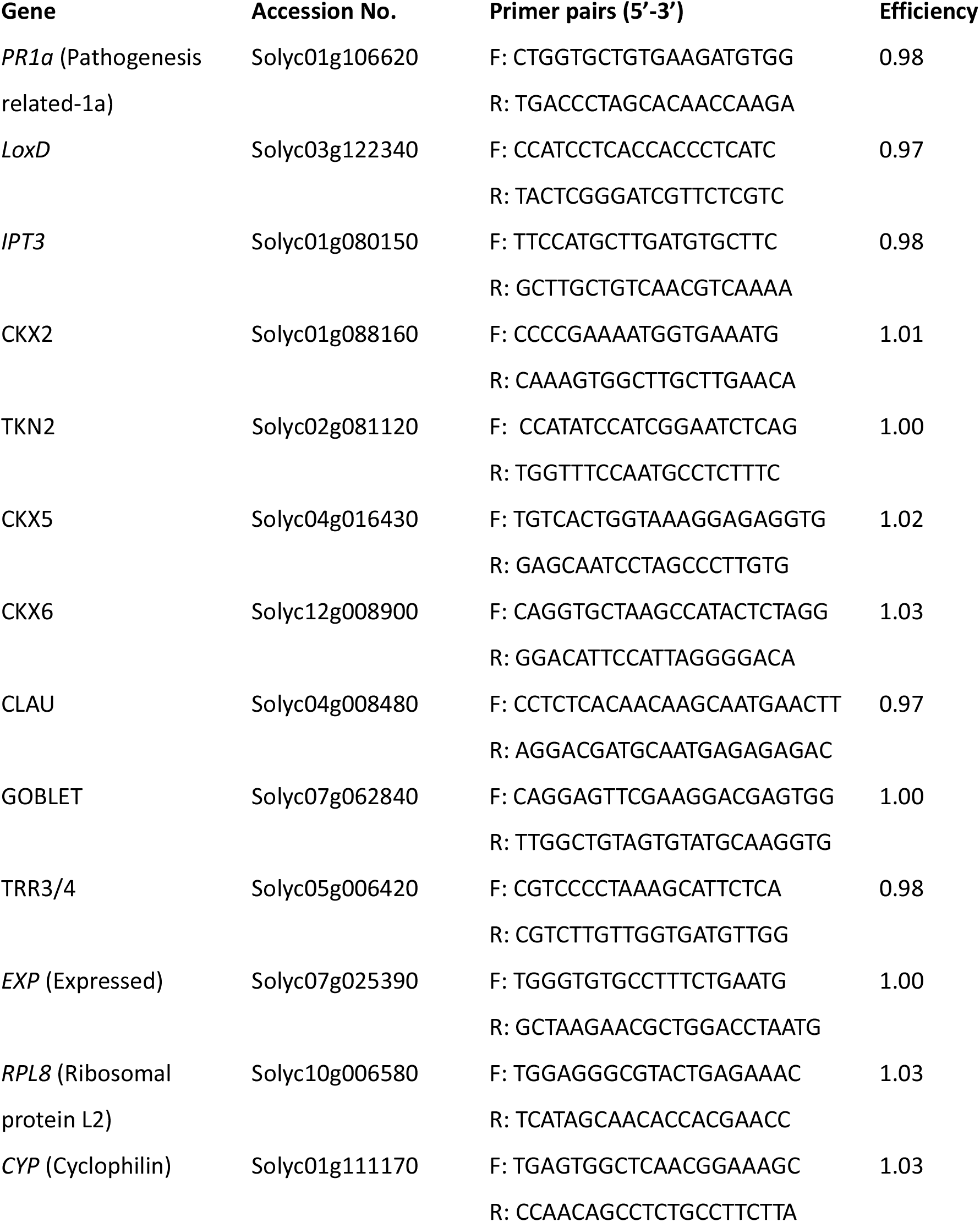
Primers used for qRT-PCR.

